# Building clone-consistent ecosystem models

**DOI:** 10.1101/724898

**Authors:** Gerrit Ansmann, Tobias Bollenbach

## Abstract

Many ecological studies employ general models that can feature an arbitrary number of populations. A critical requirement imposed on such models is *clone consistency*: If the individuals from two populations are indistinguishable, joining these populations into one shall not affect the outcome of the model. Otherwise a model produces different outcomes for the same scenario. Using functional analysis, we comprehensively characterize all clone-consistent models: We prove that they are necessarily composed from basic building blocks, namely linear combinations of parameters and abundances. These strong constraints enable a straightforward validation of model consistency or reveal implicit assumptions required to achieve it. We show that such implicit assumptions can considerably limit the applicability of models and the generality of results obtained with them. Moreover, our insights facilitate building new clone-consistent models, which we illustrate for a data-driven model of microbial communities. Finally, our insights point to new relevant forms of general models for theoretical ecology. Our framework thus provides a systematic way of comprehending ecological models, which can guide a wide range of studies.

## I. INTRODUCTION

Many theoretical and semi-empirical studies of ecological communities employ general models that are not specific to a given community, but can incorporate an arbitrary number of populations with different properties [1–4]. In most such models, the equations governing each population have the same shape, and the species of a population only manifests in the values of the associated parameters. These parameters may describe the properties of a single population, the interaction of two populations, or higher-order interactions, i.e. effects involving three or more populations [5, 6]. Interaction parameters are often chosen randomly [7–13] or determined from experiment [14–17].

Developing such models is one of the challenges of modern ecology, in particular when incorporating empirical data [18]. For example, recent advances in automating experiments have enabled measuring interaction parameters for richer communities [16,19–21], interactions characterized by more than one parameter per pair of populations [16], or higher-order interactions [15, 22]. These new experimental scenarios often call for new ecological models that can incorporate the respective data. One reason for this is that there is no single answer as to how multi-parameter or higher-order interactions should be measured [3, 6, 16, 20, 23]. Another driver of new modelling approaches is growing computational power [18, 24], which allows to investigate increasingly general and complex models [12, 25].

To improve the modeling process, several collections of criteria capturing consistency were suggested [26–34]. While many of these are specific to the ecological scenario under consideration, e.g., predation, the following invariance is a recurring theme [28–38]: If two populations have identical parameter values, they contain identical individuals (clones) within the model. Thus, the outcome of the model must only depend on the total abundance of these two populations, and not on how the clones are assigned to them. We denote this criterion as *clone consistency*. Similar criteria for models have been named *invariance under relabeling* [32] or *under identification/aggregation of identical species* [29–31] as well as *“common-sense” criterion* [34, 36, 38]. Further, it is often required that joining two populations of identical individuals does not affect diversity measures and other ecological observables [39–41]; Ref. 42 introduced this concept under the name *twin property*. Finally, in the analysis and modelling of food webs, this issue is essentially circumvented by considering *trophic species*, which aggregate species with identical predators or prey – a controversial approach [43–47].

To provide an instructive example for clone inconsistency, we compare two simulations of a predator-prey scenario using the same model [48] (chosen here exclusively for its simplicity):

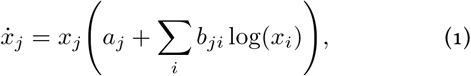

where *x_j_* is the abundance of population *j*, and *a* and *b* are parameters governing the properties and interaction of the populations. In the first simulation (solid lines in Fig. 1), we chose:

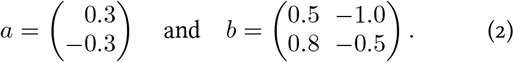

**Figure 1.**
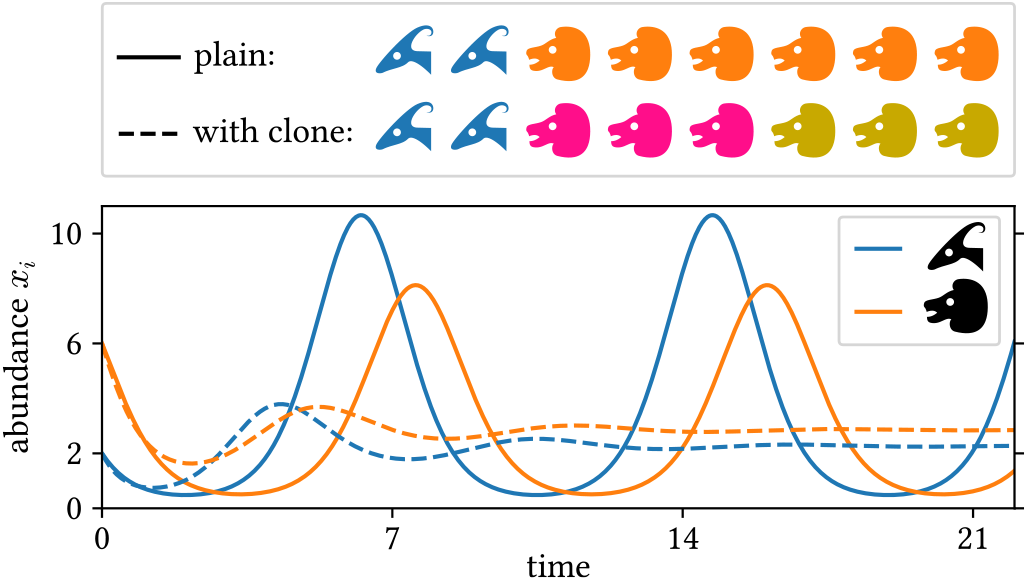
Example of clone inconsistency: two simulations of a simple predator–prey scenario using the same model (see text for details). Solid lines: simulation using one population for prey (blue, antelopes) and predators (orange, lions) each. The initial abundances are *x*_1_(0) = 2 and *x*_2_(0) = 6 (animal heads in the top legend). Dashed lines: same, but with two identical predator sub-populations (pink and ocher) with half the initial abundance; the abundance shown for the predators is the sum over the two sub-populations.

Population 1 is the prey; Population 2 is the predators. In the second simulation (dashed lines in Fig. 1), we split the predator population into two sub-populations (2 and 3) with identical properties and half the initial abundance. Allegorically, we paint half the predators in a different color. In numbers:

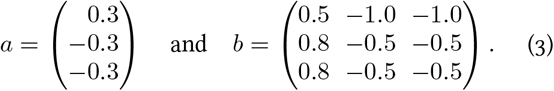

Although these two simulations describe the same situation, their outcomes differ strongly (Fig. 1): Not only does the amplitude of the predator and prey abundances change, but the type of population dynamics changes from an oscillation to a simple convergence on a fixed point. These incompatible results show that the model used in these simulations suffers from a fundamental inconsistency. In this work, we show that this is an inevitable consequence of the use of the logarithm and that log(*y*) + log(*z*) ≠ log(*y* + *z*).

While splitting populations is an illustrative thought experiment, its implications reach farther for at least three reasons: First, in actual modelling we can encounter the inverse situation, i.e., two populations with identical properties. Second, if there are problems when two populations have absolutely identical properties, there will also be problems when they have similar properties since models, like nature, are continuous. Third, problems can already arise if two populations are similar in one aspect that is relevant to the model. In a variation of the above example, if the members of two populations of predators prey with a similar rate on a given focal species, assigning individual predators to the other population should not disproportionately affect the total predation rate.

Despite its simplicity, ensuring clone consistency directly can be tedious as it requires finding a counter-example or performing a model-specific proof. For example, Ref. 33 spent several pages of calculations on checking a weaker criterion for a handful of models. Several proposed models [3, 16, 48–60] are not clone-consistent (see Sec. IV for details). Here we mathematically encode clone consistency together with a few other basic criteria. Applying tools from functional analysis, we expose the consequences of these criteria and derive a framework that enables straightforward consistency checks for any given model. Our criteria further yield a recipe for building consistent models from simple building blocks. We demonstrate these features by applying our approach to a recent model for microbial communities. Moreover, we discuss circumstances under which our criteria can be relaxed; even then, they can reveal implicit assumptions of a model and guide modelers. Finally, we point out general implications of our theoretical results for modeling studies.

#### Box 1. Mathematical Notation

**lowercase italic letters:** numbers or parameter configurations (tuples of numbers);
**lowercase Greek letters:** functions;
**boldface letters:** vectors or similar;
**uppercase letters:** sets of respective contents;
*n*: the number of populations;
*m*: the number of parameters per population;
ℝ_+_: the non-negative real numbers;
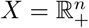: the space of all possible population abundances;
*A* = ℝ^*n*×*m*^: the space of all possible parameter configurations of these populations;
*X* × *A*: the domain of impact functions;
**x** = (*x*_1_,…,*x_n_*) ∈ *X*: an arbitrary first argument of an impact function (abundances), where *x_i_* is the abundance of population *i*;
**a** = (*a*_1_,…, *a_n_*) ∈ *A*: an arbitrary second argument of an impact function (parameters), where *a_i_* ∈ ℝ^*m*^ are the parameter values that describe population *i*;
**non-italic sans-serif letters:** modifications of specific components of arguments of an impact function (similar to named arguments in many programming languages). For example: *φ*(**x, a**, x_2_ = *y*) denotes *ϕ*((*x*_1_, *y*, *x*_3_,…, *x_n_*), (*a*_1_,…, *a_n_*)).
Here the arguments of the function *ϕ* are **x** and **a** except for the abundance of the second population (x_2_) being changed to *y*.
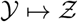: the function that maps 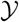 to 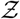 (anonymous function).
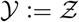: 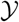 is defined as 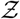.

## II. WHAT MODELS ARE CLONE-CONSISTENT?

### A. Defining and Constraining Impact Functions

To investigate the consequences of consistency requirements on entire ecosystem models, we introduce *impact functions:* These functions describe the impact of a community on a species, on a resource, or on any other relevant feature of the ecosystem that is captured in a model. Features and phenomena described by impact functions include:

- the effective growth rate of a given species,
- the remaining size of a niche,
- the availability of a nutrient,
- the rate of predation,
- reproductive services, e.g. pollination,
- the amount of crowding,
- general interaction terms, e.g. the sum in the Volterra model [1].

The arguments of impact functions are the abundances of all populations in the ecosystem **x** = (*x*_1_, *x*_2_,..., *x_n_*) and parameters **a** = (*a*_1_, *a*_2_,..., *a_n_*) that quantify the impact of the populations. Often the impact of a population *i* is described by a single number. Yet our results also hold for the more general case that *m* parameters per population are required, i.e., *a_i_* = (*a*_*i*1_, *a*_*i*2_, …, *a_im_*)∈ ℝ^*m*^.

A prominent example of an impact function is:

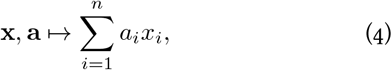

where 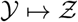 denotes the function that maps 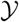 to 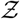. This is employed for the interaction term in the Volterra model [1] amongst others. In general, impact functions can take many forms and are a central ingredient of ecological models.

We require impact functions to fulfill the following basic criteria (illustrated in Fig. 2; see Appendix A for mathematical formulations):

**FIG. 2.**
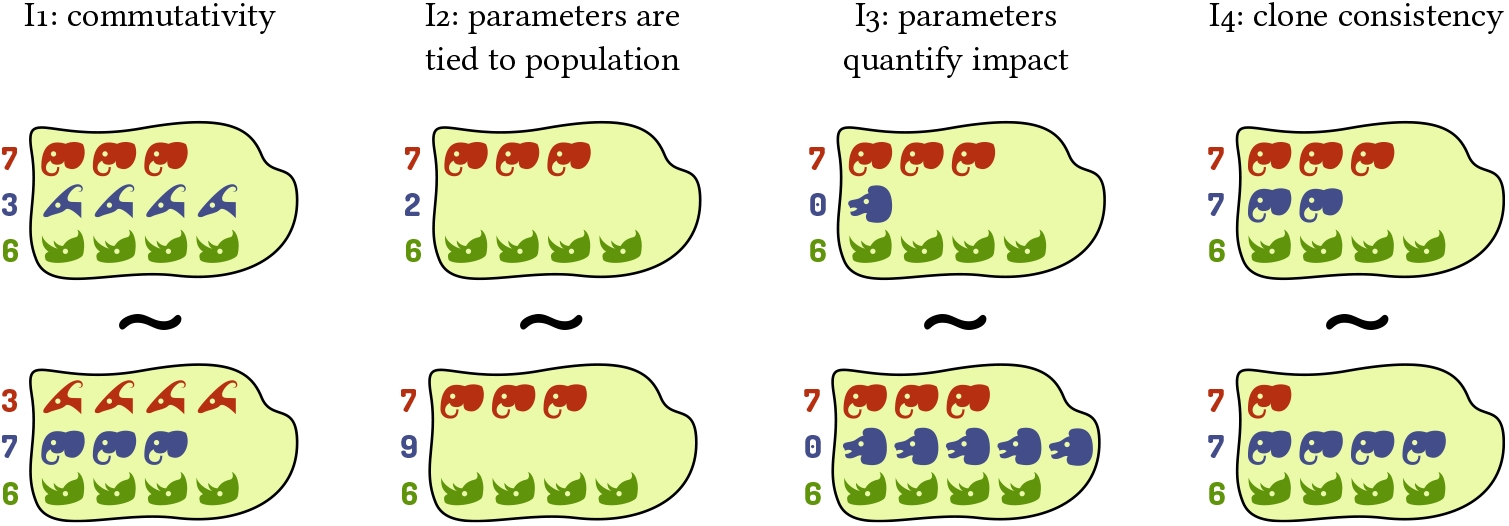
Criteria for impact functions exemplified for the grazing impact of animal populations. Each island represents a community. Each row and color represents one population in the model, with animal heads representing individuals. Numbers on the left represent parameters governing the respective population (grazing rate in the example), and head shapes indicate whether populations have identical properties as per these parameters. The similar sign (~) indicates that two communities are equivalent as arguments of an impact function, i.e., they should yield the same result (total amount of grazing in the example).

**I_1_:** Commutativity: The properties and idiosyncrasies of a given population are exclusively captured by its associated parameters, as opposed to dedicated mathematical terms in the function. This is equivalent to pairs of abundances and parameters ((*x_i_,a_i_*)) being interchangeable as arguments of the impact function.

**I_2_:** When a population is absent, its associated parameters have no effect on the value of the impact function.

**I_3_:** When the parameter(s) associated with a given population are zero, that population’s abundance has no effect on the value of the impact function.

**I_4_:** Clone consistency: If two (or more) populations have identical parameters, the value of the impact function must only depend on their summed abundance and not on its distribution among the two populations. As our scope here is impact functions, the populations do not need to be identical in all respects, but only in the parameter(s) used by the respective impact function.

Note that it is often reasonable to choose a considerable portion of parameters to be zero. For example, if our impact function describes predation loss of a given focal species, we would choose *a_i_* =0 for all populations *i* that do not prey upon the focal species. In the following we call a function *impact function* if and only if it satisfies these criteria.

### B. The Shape of Impact Functions

While the criteria I_1_–I_4_ for impact functions are conceptually simple, it is not straightforward to test directly if a given function complies with them or to devise a new function that does. To address this issue, we investigated the functional shape of impact functions and found a set Ω of *basic impact functions*, which can serve as building blocks for ecosystem models.

These basic impact functions are linear combinations of abundances and parameters (potentially transformed), i.e., all functions of the shape:

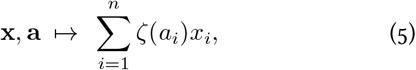

where ζ : ℝ → ℝ is an arbitrary parameter transformation with ζ(0) = 0. Often, ζ is the identity function (id), simplifying Eq. 5 to Eq. 4. Using methods from functional analysis, we mathematically proved that any impact function can be built from these basic impact functions via addition, multiplication, function composition, and similar operations (see Appendix A). Conversely, everything built from those elements or other impact functions is again an impact function. Formally, the general functional shape of impact functions is:

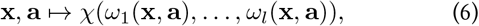

where *ω*_1_,..., *ω_l_* ∈ Ω are basic impact functions as per Eq. 5 and *χ*: ℝ^*l*^ → ℝ is an arbitrary function combining their results, such as a product, sum, or a more complex function.

To illustrate the composition of impact functions from these building blocks, we consider the case of a single population of flowering plants that may be both pollinated and grazed upon by several insect populations (see, e.g., Ref. 61) as a toy example (also see Fig. 3, bottom). We use a function *ρ*_graz_ to describe the rate at which insects (and their larvae) graze on the plants:

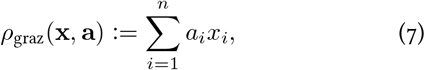

where *a_i_* is the grazing rate of insect population *i*. The function *ρ*_graz_ has the shape of Eq. 5 (with *ζ* = id) and therefore it is a (basic) impact function. We use a function *ρ*_poll_ to capture the rate at which insects fertilize flowers:

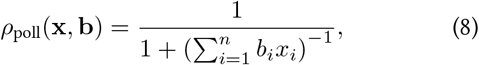

where *b_i_* is the contribution to pollination provided by insect population *i*. A Holling type-II response here ensures that the fertilization rate saturates at 1. Again, we can see that this is an impact function as it is built via composing 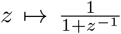 and the basic impact function 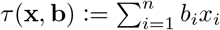. Finally, we combine the two impact functions to a function *ϕ* that describes the relative change of the plant population due to insect influence:

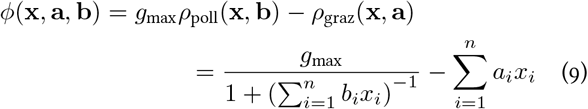

where *g*_max_ is the growth rate of the plant population in the absence of death and with maximum fertilization. As *ϕ* is built from impact functions, it is an impact function itself. We can also see directly that *ϕ* complies with the general shape of an impact function (Eq. 6) by choosing 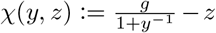, *ω*_1_ = *τ*, and *ω*_2_ = *ρ*_graz_. As a result, we can be certain that the resulting model satisfies the fundamental consistency criteria. Similarly, the basic building blocks we identified enable constructing consistent models of any other ecosystem.

**FIG. 3.**
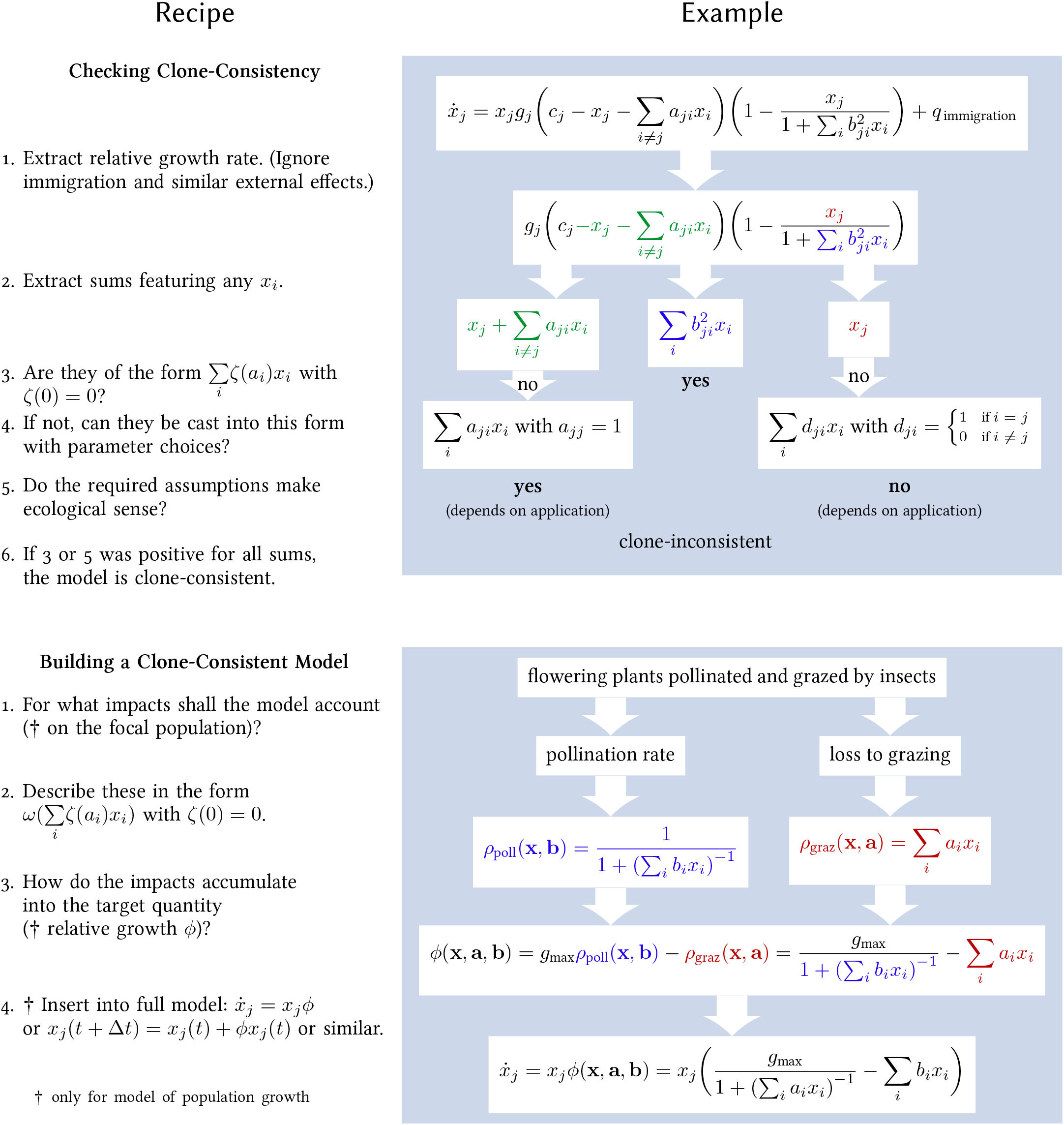
Recipes for building and validating models using our framework. In general *x_i_* denotes the abundance of population *i*, and *a_i_, b_i_, c_i_*, and *d_i_* are parameters describing its impact. The example for checking is based upon Eq. 20 and tailored for covering relevant cases. The example for building extends the one from Sec. II B. For both examples, some specifics are discussed in more detail in the main text.

### C. Assessing and Building Models

Our treatment of impact functions allows us to ensure that we are clone-consistent when modeling the effects **of** a community. We now additionally consider clone consistency when modeling the effects **on** the growth of a population. Specifically, we require that the total growth of two populations of identical individuals must be the same as if those two populations were joined in the model.

We consider models of the forms:

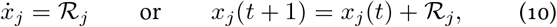

where the right-hand side 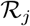 describes the change of abundance of population *j* due to the internal dynamics of the ecosystem. For simplicity, we omit the dependencies of 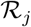 and ignore immigration and other external effects, but adding them to such a model is straightforward.

We proved that 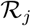 must have the shape (see Appendix C):

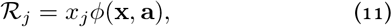

where *ϕ* is an impact function. This means that all dependencies of 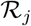 on **x** must either happen within an impact function or in the form of a single factor *x_j_*. Intuitively, each individual multiplies with a rate that is the result of all impacts it experiences in the ecosystem (*ϕ*) – these impacts include interactions between individuals of the same population, e.g. due to crowding.

This insight provides an easy way to verify if models comply with our consistency criteria. We simply need to check if they have the shape of Eq. 11. To verify in turn if *ϕ* is an impact function, we can look for terms of the shape of Eq. 5. For instance, the Volterra model [1] can be rewritten as:

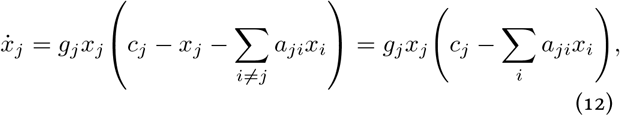

with *a_jj_* = −1, which reflects that a population maximally competes with itself. With this, the Volterra model is clearly built from a linear combination and a factor *x_j_* and we can thus be sure that the model is clone-consistent. We can now have a another look at our introductory example (Eq. 1): As the interaction term features logarithms of abundances, it does not comply with the shape of Eq. 5 and thus the model violates our consistency criteria. Hence the observed clone inconsistency (Fig. 1) is inevitable. We summarize the recipe for checking a model and provide a more extensive example in the top of Fig. 3.

This recipe can also be inverted to build a clone-consistent model. In the example from Sec. II B, we can directly insert into Eq. 11 and obtain a model for the change of a plant population in light of pollination and grazing (see Fig. 3 bottom). Note that if we stringently apply our framework here, plants are distinguished from insects only by having a grazing and pollination rate of zero.

Our framework can also be applied to experiments that do not assess the details of ecological interactions (nutrients, toxins, etc.) but only aggregated phenomenological observables such as the carrying capacity of a population in the presence of another. This is typical for high-throughput experiments assessing the pair-wise interactions of microbial communities [15,16,19–22]. For tailoring a model to the data from such experiments, we propose to make a general ansatz using one basic impact function for each of the *m* experimentally determined interaction parameters, for example:

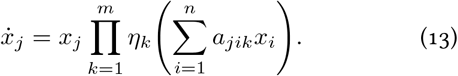

The parameters and functions (*a_jik_* and *η_k_*) can then be determined using:

- the requirement that the observed experimental scenarios should be reproduced by the model,
- ecological assumptions and facts about the scenario, e.g., that the predation rate should increase with the abundance of predators, or
- assumptions of simplicity (Occam’s razor).

Using fewer building blocks would imply that experimentally determined parameters are not used, i.e. available information on the system would be ignored; more building blocks would usually result in models that are overly complex in light of our limited knowledge about the system. We refrained from transforming the parameters in Eq. 13 (corresponding to *ζ* = id in Eq. 5), since this would not affect the final model. Finally, for some applications, a sum or more complex way to combine the basic impact functions may be appropriate (as opposed to the product used in Eq. 13). In Sec. III B, we provide an example for this approach.

### D. Higher-Order Interactions

Our approach is readily extended to models describing second- or higher-order interactions. For this, one simply has to consider parameters that are associated with more than one population. It is then convenient to use other basic building blocks (instead of Eq. 5), for example for second-order inter-actions:

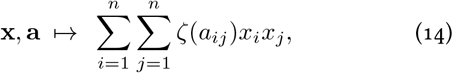

which is an impact function independent of whether *a_ij_* is considered a parameter associated with population *i* or with population *j*. Analogous building blocks exist for higher interaction orders. Such building blocks are featured in existing models that capture higher-order interactions [12, 13, 17].

## III. CASE STUDY: SEMI-EMPIRICAL MODELS FOR MICROBIAL COMMUNITIES

As an instructive example, we apply the impact-function framework to describe the dynamics of a microbial community for which multiple ecological interaction parameters were recently measured experimentally. Ref. 16 (co-authored by one of us) used a high-throughput approach to systematically measure ecological interactions in microbial communities consisting of strains isolated from polymicrobial urinary-tract infections (UTI). For each strain, the exponential growth rate *g_j_* and the carrying capacity *c_j_* in isolation were measured (Fig. 4, left). For convenience, abundances of each strain *j* were normalized such that *c_j_* = 1. Furthermore, for each strain *k*, a medium partially conditioned by that strain was produced (it contains a fraction *v* of the supernatant). In each such partially conditioned medium, the conditioned growth rate *g_jk_* and carrying capacity *c_jk_* of each strain *j* were measured to quantify how strain *k* affects strain *j* (Fig. 4, right). Creating a model from this data is particularly challenging since it has to include two experimental interaction parameters: growth rate and carrying capacity in conditioned medium.

**FIG. 4.**
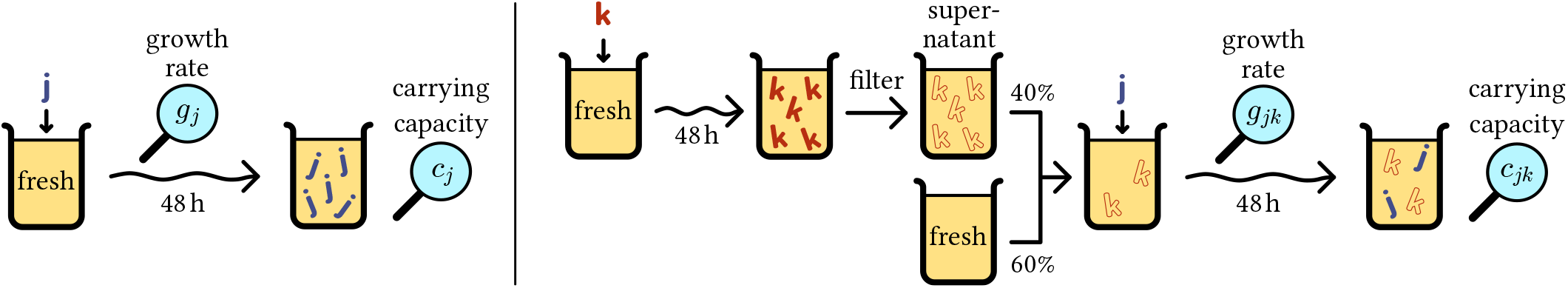
Acquisition of data used in our case study. Measurement of growth characteristics (left) and pairwise interactions (right) of bacterial strains isolated from urinary-tract infections by Ref. 16. Left: Each strain *j* was cultivated for 48 h in artificial urine. Solid bold letters represent individuals of the respective strain. The exponential growth rate *g_j_* as well as the carrying capacity *c_j_* (named *yield* in Ref. 16) were experimentally determined via optical densities. Right: For each strain *k*, a conditioned medium was produced by letting the strain grow for 48 h, mechanically removing the bacteria to obtain a *supernatant*, and mixing the result with fresh medium in a ratio of *v* ≔ 0.4. Outline letters (“footprints”) indicate to what extent the respective culture consists of supernatant. In each such medium, each strain *j* was cultivated, and the conditioned growth rate *g_jk_* and carrying capacity *c_jk_* were determined as above.

For modelling this system, it can be assumed that the abundance of a population also represents its footprint, i.e. the nutrients, toxins, and other relevant substances produced or depleted by that population. The basis for this simplifying assumption is that populations decline in a relevant magnitude only due to dilution of the entire system, which equally affects the footprint. With this assumption, we can treat the medium partially conditioned by strain *k* as an ecosystem where the abundance of that strain is fixed to the corresponding fraction of its carrying capacity (*x_k_* = *v_c_k__* = *v*).

The general form of an ordinary differential equation describing the (normalized) abundance *x_j_* of population *j* in this system is 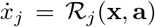. A sensible criterion for such models is that they should reproduce the observed growth rates and carrying capacities for all situations that were experimentally investigated. For instance, in the absence of other strains, the initial exponential growth rate of strain *j* in the model should be equal to the experimentally observed exponential growth rate *g_j_*:

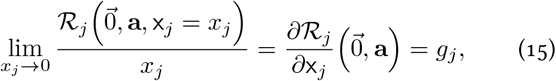

where the argument 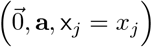 of 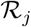 denotes that the abundance of population *j* is *x_j_* and all other abundances are zero. Similarly, we can deduce three other criteria, resulting in one per experimental interaction parameter in total:

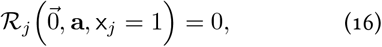

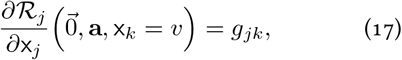

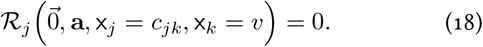

### A. Existing Model

Ref. 16 proposed a model for communities consisting of such strains based on Verhulst’s logistic model for one population [62]:

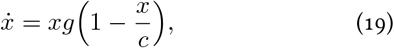

where both the growth rate *g* and the carrying capacity *c* (normalized to 1 here) are modified by interaction terms incorporating the experimentally obtained parameters:

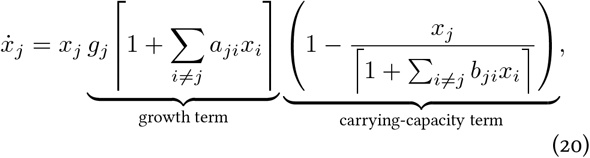

with ⌈*z*⌉ ≔ max(0, *z*), 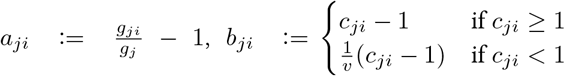, and the remaining symbols as in Fig. 4. Looking at this model through the lens provided by our framework, both the carrying-capacity and the growth term should be impact functions, with *a* and *b* being the parameters quantifying these impacts. However, it is clear that neither term is built from linear combinations (with complete sums), matching the shape of Eq. 5. As the model clearly satisfies Criteria I_1_–I_3_, it therefore must violate I_4_ and be clone-inconsistent which can indeed be shown explicitly (Appendix D). We can complete the sums with appropriate choices of *a_jj_* and *b_jj_*, and we can expand the solitary *x_j_* in the numerator to:

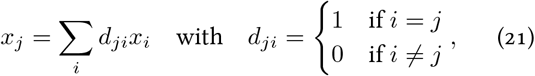

where the new parameter *d_ji_* can be interpreted as quantifying the extent to which population *i* occupies the niche of population *j*. However, this expansion implicitly assumes that the strains have non-overlapping niches, which is not justified for this study, as it features many communities containing two strains of the same genus or even species. This interpretation of *d_ji_* would also ignore that the conditioned carrying capacities *c_ji_* capture niche overlap.

The fixed points of this model are characterized by:

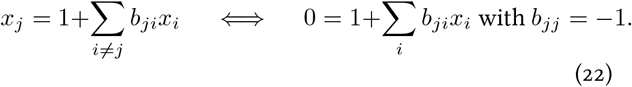

Interestingly, they thus are clone-consistent (also see Fig. 5). Most conclusions of Ref. 16 are based on these fixed points and thus unaffected by the clone inconsistency of the model. However, clone inconsistency affects the transient dynamics (see also Fig. 5), which is relevant as these communities are subject to frequent dilutions (due to bladder voiding) which can happen long before the system has equilibrated.

**FIG. 5.**
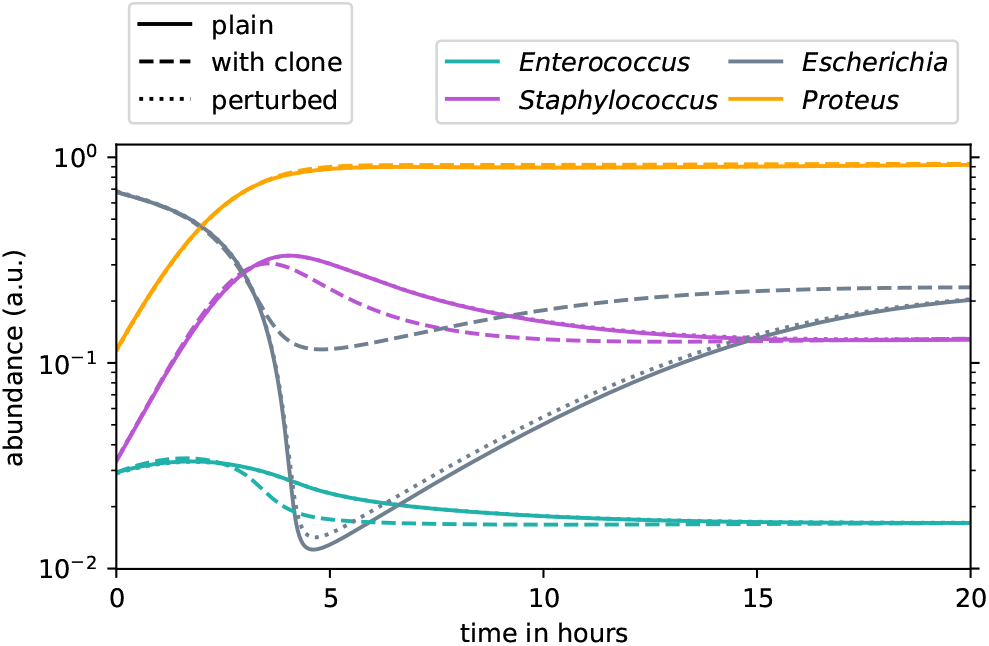
Simulation of a UTI community (Community 3 from Ref. 16) using Eq. 20. Solid lines: plain simulation; each strain is represented by one population. Dashed lines: the same, except that there are two *Enterococcus* populations with identical parameters and half the initial abundance each. The abundance shown for *Enterococcus* is summed over these two populations. Dotted lines: like plain simulation, except that the initial abundances were perturbed by 1 % in a random direction. (A dotted line may be mostly covered by the respective solid line.) This demonstrates that the difference between the former two simulations is not caused by numerical noise and sensitivity to initial conditions. See Appendix F for details of the simulation.

### B. New Model

We apply our framework to construct a new population dynamics model for this scenario based on the same experimental data. Since two experimental interaction parameters are available, we make an ansatz using two basic impact functions (see Sec. II C). As one of the parameters and thus one of the impact functions reflects the carrying capacity, we choose to multiply (and not add) the impact functions to ensure that an impact function can single-handedly reduce the growth to zero. Our ansatz is thus Eq. 13 with *m* = 2:

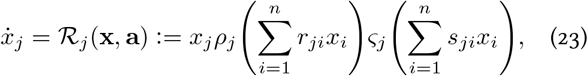

where *a*_*j*1_ ≔ *r_j_* and *a*_*j*_2__ ≔ *s_j_*.

Inserting this ansatz into Eqs. 15–18 and making a few choices that do not affect generality already yields strong constraints on the functions *ρ_j_* and *ζ_j_* and on how the parameters *r_jk_* and *s_jk_* relate to these and the experimental parameters *g_j_*, *g_jk_* and *c_jk_*, namely (see Appendix E): 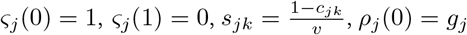, and 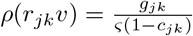. By further making simple choices for *ρ_j_* and *ς_j_* within these constraints and accounting for singularities and discontinuities (see Appendix E), we arrive at the following model (with ⌈*z*⌉ ≔ max(0, *z*)):

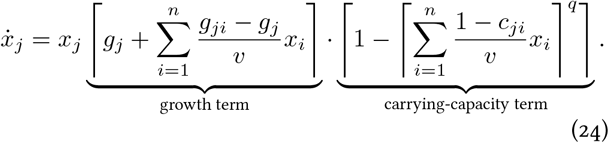

Like the existing model (Eq. 20), this can be understood as an expansion of Verhulst’s logistic model with the following differences: First and foremost, the carrying-capacity term is simplified to achieve clone consistency: Interactions affecting the carrying capacity and its utilization by population *j* are now captured in a single sum. Second, the additional parameter *q* governs how abruptly the saturation effect kicks in. Third, the dilution factor *v* is included consistently without case distinctions. Fourth, as per the initial assumptions, populations cannot decline anymore (unless dilution is added to the model). The particular shape of this model illustrates how challenging it can be to write down clone-consistent models from scratch without using the framework presented here.

We find that this model performs at least as well as the previous one (Eq. 20) when it comes to predicting a small experimental dataset (Appendix F).

## IV. IMPLICATIONS

### A. Checking Clone Consistency to Reveal Implicit Assumptions

While many popular models such as most variations of the Volterra model [1] comply with our criteria, others do not (e.g. the models from Ref. 48, 50, 53, 54, 57, and 58, Eqs. 9 and 10 in Ref. 49, Eqs. 1.28-1.30 and 1.50 in Ref. 51, Eqs. 11 and 12 in Ref. 52, Eq. 5 in Ref. 55, the NFR model in Ref. 56, Eq. 3 in Ref. 59, Figs. 3b and c in Ref. 3, the UIM and IIM model in Ref. 60, Eq. 20). However, as we will elaborate below, this does not necessarily mean that these models should be dismissed outright.

As we illustrated in Fig. 3 and Eq. 21, it is possible to transform every model into a clone-consistent one – if one assumes that the right parameter values are zero. For instance, suppose that *u* – *x_j_* quantifies the size of the unoccupied portion of the niche of population *j*. We can extend the latter term to an impact function:

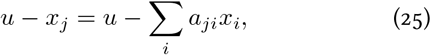

with *a_ii_* = 1 and *a_ji_* = 0 for *j* ≠ *i*. Here *a_ji_* describes the extent to which population *i* occupies the niche of population *j*, and thus *a_ji_* = 0 for *j* ≠ *i* implies that population *j* exclusively occupies its niche. Whether this assumption of an exclusive niche can be justified depends on the application. The strength of our framework is to systemically point to such (usually implicit) assumptions and prompt an explicit justification or an improvement of the model if the assumptions are not justified.

In a specific example, the unique-interactions model (UIM) from Ref. 60 features three interaction terms. Two of these terms describe the influence of mutualistic or exploitative interactions, respectively, on a focal population *j* as:

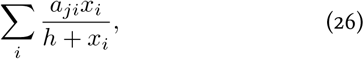

where *a_ji_* describes the strength of the respective interaction (and is zero when it is absent) and *h* is the half-saturation constant. When taken as they are, these terms are not impact functions and thus clone-inconsistent. However, an eponymous assumption of this model is that mutualistic and exploitative interactions are unique for a given focal population, which we consider a valid approximation for the purposes of that work. Per this assumption, each summand of Eq. 26 represents a separate interaction mechanism, such as a specific resource, service, or mode of predation, and only population *i* affects the focal population *j* via this mechanism. Thus, in our framework, each summand can be expanded to a single impact function, with all but one parameter value being zero. By contrast the third term in the model of Ref. 60 describing competitive interactions is clone-consistent, and here unique interactions are neither assumed nor would this be justified.

Many of the aforementioned potentially clone-inconsistent studies and modelling studies primarily make statements about how model properties (such as types of interaction) affect population dynamics. If these findings are subject to strong assumptions, this considerably diminishes their generality, relevance, and applicability. Moreover, if substantial assumptions are implicit, this increases the risk that the study is being misinterpreted and misapplied by others.

For some features of populations, it is not feasible to encode them in numerical parameters – which we assumed as given so far. For example, compatibility for sexual reproduction is tedious to capture in a parameter and a more useful approach is to assume that populations do not interbreed (i.e., contain species as defined by Mayr). In this case, splitting a population in two equal parts also halves the availability of partners for sexual reproduction. Thus, this availability should not be described by a clone-consistent impact function.

Finally, we emphasize that our approach extends to diverse types of models. In particular, it is not restricted to models employing ordinary differential equations, but can also be applied to models with noise, time delays, or discrete time steps, e.g., one per generation. Also, higher-order interactions are covered by our framework. Moreover, while we mainly used impact functions to describe the impact of a community on a population, both the targets and the source can be other entities, e.g. the availability of a resource, the concentration of a toxin, or an aggregated observable such as the albedo of foliage or the pH value of a medium. A common case is the impact of the community on a resource within a consumer–resource model [63, 64]. Going beyond modelling, impact functions targetting observables ties into requirements for ecological observables like diversity to be clone-consistent [39–41].

### B. General Implications for Model Design

While each specific case of unjustified clone inconsistency reflects a biological shortcoming, it is not possible to make general qualitative or quantitative statements about the causes or consequences of clone inconsistency. This is because clone inconsistency is inevitably intertwined with the fabric of the model, and thus we cannot study its effect in isolation. Moreover, clone-inconsistent models are diverse for the same reason that non-linear functions are. For example, it makes a difference whether joining identical populations increases or decreases some impact (which would be unchanged in a clone-consistent model). However, there are some insights to be gained from interpreting each impact function as one mechanism of ecological interaction:

Many pure modeling studies use a model of the general shape [12, 56, 59]:

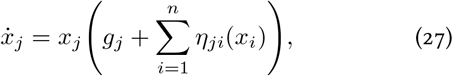

where *g_j_* is the unperturbed growth rate of population *j*. If all *η_ji_* are linear, this kind of model employs a single basic impact function and requires no further assumptions. If, on the other hand, the *η_ji_* are non-linear, this can only be justified within our framework as follows: Each summand in Eq. 27 corresponds to one interaction mechanism, with which population *i* uniquely affects population *j*. In this case, each summand would be expanded to an impact function for each of which all but one parameter are zero, which makes for a total of *n* basic impact functions. An interesting implication is that models of the above shape can either feature 1 or *n* basic impact functions (depending on whether *η_ji_* is linear), but they cannot capture the middle ground in between. As each basic impact function can be associated with one interaction mechanism, this limitation is relevant beyond the consistency issues addressed by our framework.

To fill this gap, our framework suggests an alternative shape for general ecosystem models:

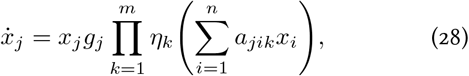

where *m* is the number of impact functions (compare to Eq. 13). Each factor in Eq. 28 corresponds to one interaction mechanism, which can involve multiple populations. To reflect that most populations do not participate in most given interaction mechanisms, we suggest that the interaction coefficients (*a_ijk_*) are sampled from a distribution that contains a considerable portion of zeros. The above general model can inform studies employing random interaction parameters [9–13], general modeling [25], or machine learning by narrowing down or expanding the space of possible models taken into consideration.

We also note the conceptual parallels of the framework presented here to two pharmacological approaches to describe the combined effect of two drugs [65,66]: One is *Loewe additivity*, which is based on arguments similar to clone consistency and holds if the two drugs affect the same component of the cell. The other is *Bliss independence*, which violates clone consistency at first glance and holds if the two drugs affect different components of the cell. In our framework, drugs that target the same cell component correspond to using the same interaction mechanism and thus would be captured by the same basic impact function. The effect of a complex drug cocktail could be captured by several Bliss-independent basic impact functions, each of which comprises a series of Loewe-additive components.

## V. CONCLUSION

We introduced a framework for building ecosystem models centered around impact functions as building blocks. This framework is aimed at ensuring the clone consistency of models and thus constrains the possible choices of models. While at first this may seem like at a burden, we expect that it rather eases the modelling process as it guides modelers when choosing from the (still infinitely many) clone-consistent models. Our framework further prompts relevant questions about the assumptions that enter a model. On the other hand, the absence of impact functions in a model exposes that it is clone-inconsistent or at least requires assumptions, which are often implicit and may limit the model’s generality. Our framework also informs the shape of more general models, pointing out potential new directions of research in this field and outlining the space of possible models for ecosystems. Finally, our approach could be extended to implement criteria for specific ecological scenarios such as predation [27–30, 33]. Thus, the framework presented here provides a systematic way of comprehending models and can form the backbone for a wide range of ecological modeling studies.

## Supporting information

Code for Simulations

## ACKNOWLEDGMENTS

We are grateful to H. Arndt, M. Cosentino-Lagomarsino, A. Espinosa-Cantú, U. Feudel, J. Freund, T. Gross, S. Khaiwal, Y. Mulla, G. Petrungaro, S. Vet, M. de Vos, J. Werner, and M. Zagorski for inspiring discussions or constructive comments on previous versions of the manuscript. This work was supported in part by Austrian Science Fund (FWF) standalone grant P 27201-B22 and German Research Foundation (DFG) Collaborative Research Center (SFB) 1310.

## Appendix A: The Functional Algebra of Impact Functions

Expressed in equations, our criteria for impact functions are:

**I_1_**: Commutativity:

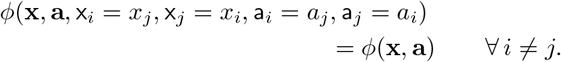
**I_2_**: When a population is absent, its associated parameters have no effect:

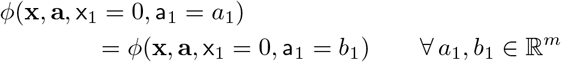
**I_3_**: When all parameters associated with a given population are zero, that population has no impact:

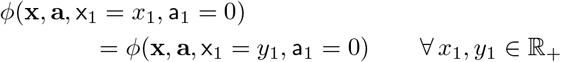

Note that the parameter value corresponding to no impact could be readily changed from zero to any other value.
**I_4_**: Clone consistency:

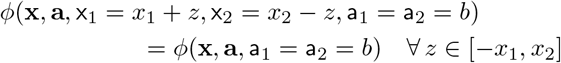

Note that through commutativity (I_1_), the other criteria apply to all populations or pairs of populations of the impact function *ϕ*, respectively (and not just to population 1 and 2). Clone consistency (I_4_) of more than two populations is covered by applying the respective criterion repeatedly.

In the terms of functional analysis, impact functions form a *functional algebra* Φ. This means that each product or sum of two impact functions is again an impact function and that each multiple of an impact function is an impact function. This algebra is also closed, which means that the limit of uniformly converging sequences of impact functions is again an impact function.

To easily build and detect impact functions, it is crucial to find a (small) set of impact functions from which all impact functions can be build, i.e., a *generating set* of Φ. Our main mathematical result is that Ξ = Λ∪Γ is such a generating set, where Γ is the set of constant functions and taking the limit of a uniformly converging sequence is considered amongst the generating operations. Λ is the set of all linear combinations of powers of parameters and abundances, i.e., functions of the shape:

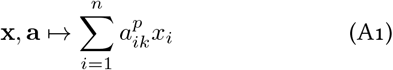

for some *p* ∈ {1,...} and for some *k* ∈ {1,..., *m*}.

We will formally prove this in Appendix B, but the essential idea is this: Our criteria (I_1_–I_4_) require an impact function to have the same value on given subsets of its domain. For Ξ to be a generating set of Φ, it must reflect this: First, Ξ must be constant on each such subset. Otherwise it would generate more than impact functions. Second and more crucially, for each pair of points that are not in the same subset, there must be a function in Ξ that differs between these points (this is called *point-separating*). Otherwise some impact functions could not be generated by Ξ. We show the latter by using our criteria (I_1_–I_4_) to systematically transform arguments of impact functions to a *canonical form*, in which populations are ordered by impact and maximally lumped together.

In application, the fact that Φ is a closed functional algebra, i.e., that limits remain within it, is relevant to capture the application of a (non-polynomial, continuous) function to the entire impact function or the parameters. We can rewrite:

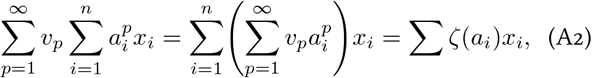

with 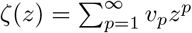. This allows us to use building blocks of the shape of Eq. 5 instead of Eq. A1. We can also rewrite:

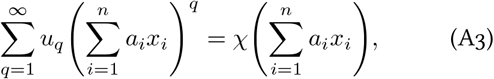

with 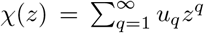. This allows for “wrapping” functions around impact functions.

## Appendix B: Proof: Ξ generates Φ

We here state and prove our main mathematical result, namely:

### Theorem 1.

*Let* 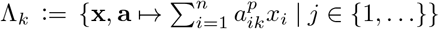 *denote the set of linear combinations of powers of values of the *k*-th parameter and abundances. Denote the set of all such functions as* 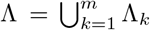. *Let* Ξ ≔ Λ ∪ Γ, *where* Γ *is the set of constant functions. Let* Ψ *be the* generated set *of* Ξ, *i.e., the smallest closed functional algebra that contains* Ξ. *Then* Ψ = Φ, *i.e*., Φ *contains all impact functions as characterized by criteria I_1_–I_4_*.

To prove it, we apply Bishop’s Theorem [67, 68], which states, when reduced to algebras of real-valued functions:

### Bishop’s Theorem

*Let Z be a compact Hausdorff space. Let* Ψ *be a closed unitial subalgebra of C*(*Z*, ℝ). *Let ϕ* ∈ *C*(*Z*, ℝ). *Suppose that ϕ*|_*S*_ *is constant for each subset S* ∈ *Z such that ψ*|_*S*_ *is constant for all ψ* ∈ Ψ. *Then ϕ* ∈ Ψ.

The requirements of Bishop’s Theorem on Ψ are fulfilled since Z can be any sufficiently large compact subset of *X* × *A* and the inclusion of Γ ensures unitiality. To show that the functional algebra Ψ contains all impact functions, we therefore need to show that for an arbitrary impact function ϕ for any 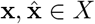 and 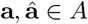:

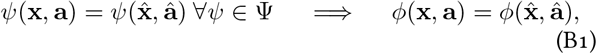

or, in the language of functional analysis, Ψ has to *point-separate*, except where no impact function point-separates. Since point-separations are unaffected by algebraic operations of functions and limits, Ψ point-separates, if and only if Ξ does. Moreover, since the functions from Γ are constant everywhere (and thus point-separate nowhere), this is equivalent to Λ point-separating. Finally, since for any *i* ≠ *j*, the functions from Λ_*i*_ are constant wherever the functions from Λ_*j*_ are not, it suffices to only consider one Λ_*i*_, i.e., scalar parameters (*m* = 1).

We prove that Λ point-separates for *m* = 1 with three lemmas, for which we transform the arguments to a canonical form (Definitions 1 – 4), in which populations are ordered by impact and maximally lumped together. We first show that if all functions from Λ have the same value for two arguments, these arguments have the same canonical form (Lemma 1). We then employ our criteria to show that no impact function will differ for two arguments that have the same canonical form (Lemma 2). Finally, we combine the first two lemmas to show that if all functions from Λ have the same value for two arguments, so do all impact functions (Lemma 3). Thus Λ point-separates.

### Definition 1.

*Let x* ∈ *X and* **a** ∈ *A. Let* 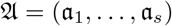 *be the ordered sequence of non-zero values of* **a** *that do not correspond to zero abundance, i.e*. 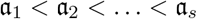 *and*:

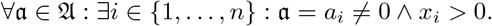

As the 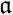 are unique and ordered, we will directly use them like indices to avoid additional levels of indexing. One can think of them as equivalence classes of parameters.

### Definition 2.

*Let* x ∈ *X and* **a** ∈ *A. For a given* 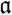, *let* 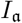 *be the set of indices where this parameter value is assumed and the corresponding abundance is not zero, i.e., the maximal set I such that* 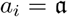 *and x_i_* > 0 *for all* 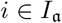. *Consequentially*, 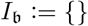 *for* 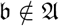.

### Definition 3.

*Let* x ∈ *X> and* **a** ∈ *A. Denote the sums of abundances for one absolute parameter value as* 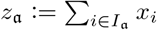.

### Lemma 1.

*Suppose* 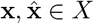 *and* 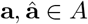 *are such that*:

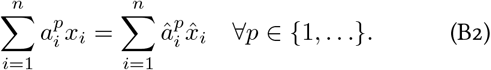

*Then:*

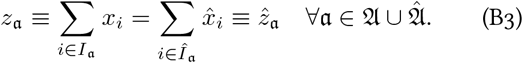

We show Eq. B3 by induction over 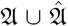 in descending order of absolute value. We first note that the lemma trivially holds for all 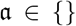. In the following we show that, if the lemma holds for all 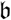 with 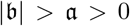, it also holds for 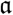 and 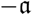. (If one of 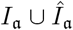 and 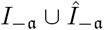 is empty, this does not affect this part of the proof.) To this end, we first show that the linear combinations must also be equal when only considering coefficients 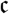 with 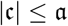 (for all *p*):

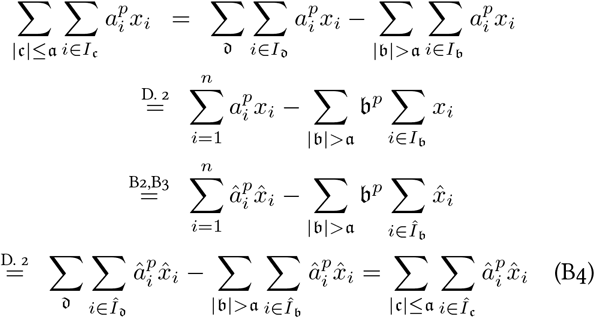

If 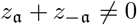 and 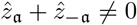 the above equality will be dominated by 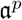 for *p* → ∞, which gives us:

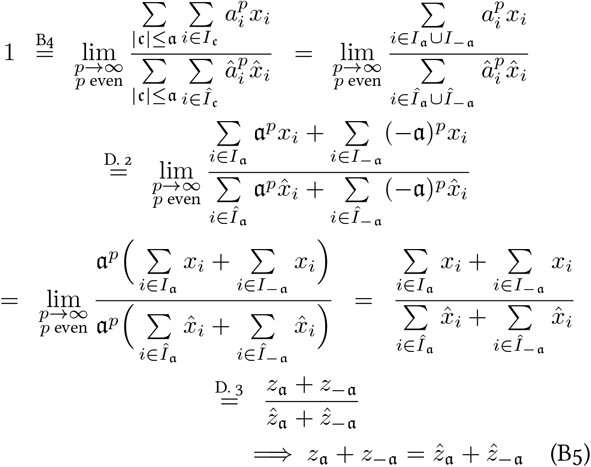

If exactly one of 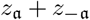 and 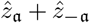 were zero, the above limit would evaluate as either 0 or ∞ instead of 1; hence this cannot be. If both are zero, Eq. B5 holds without further ado. Analogously, we obtain:

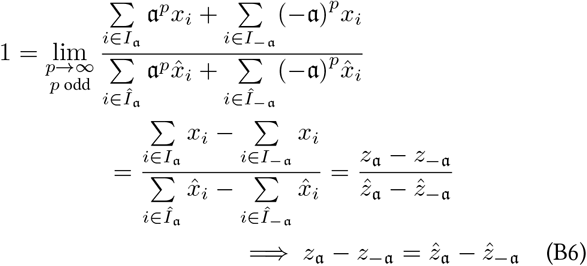

By adding and subtracting Eqs. B5 and B6, respectively, we arrive at 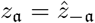 and 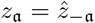.

### Definition 4.

*Define the canonical form of* **x** ∈ *X and* **a** ∈ *A as*:

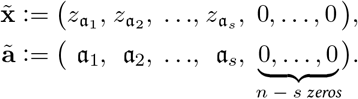

### Lemma 2.

*Let* **x** ∈ *X and* **a** ∈ *A and ϕ be an impact function. Then* 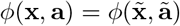.

We first transform blocks of arguments to the canonical form (with some zero arguments added if necessary) step by step, and show that the value of an impact function is not affected by these transformations. The first kind of block we consider are blocks of equal non-zero parameters and corresponding non-zero abundances, i.e., 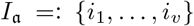 for some 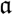. Then, for some 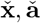:

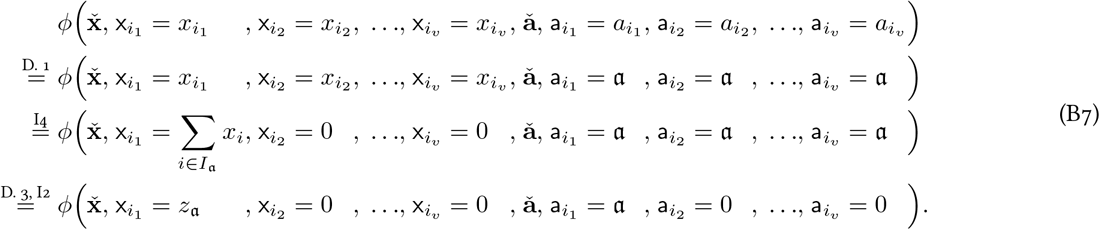

If a parameter 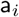 or abundance x_*i*_, respectively, is zero, we transform the single-index block {*i*} to zero (for some 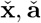):

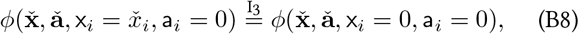

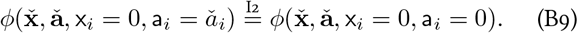

Second, after all blocks are transformed, we swap abundances and parameters in parallel to match the order in the canonical form. This does not affect the value of the impact function *ϕ* as it is commutative (I_1_).

### Lemma 3.

*Suppose* 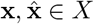 *and* 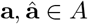 *are such that*:

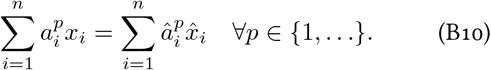

*Let ϕ be an impact function. Then* 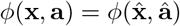.

To prove this, we only need to note how the canonical forms 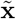 and 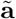 only depend on the parameters values 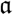 corresponding to non-zero total abundance 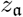 and these abundances. Those in turn are equal per Lemma 1. Thus:

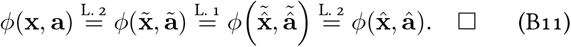

## Appendix C: Non-Impact-Function Contribution to Abundance Changes Must be Proportional

Suppose 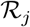 is the change of abundance of a population *j* in a model. To keep the notation simple, we assume that there is no delay, noise, or similar. We can write 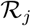 in the shape:

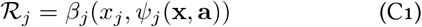

with *β_j_* : ℝ × ℝ → ℝ and *ψ* being an impact function comprising all the impacts on population *j*. If, similar to Criterion I_4_, we consider the case of two populations *j* and *k* with identical properties with abundances *y* and *z*, their total growth must be the same as if all individuals were assigned to one population:

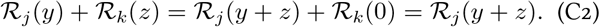

Using that *j* and *k* are identical as well as the properties of impact functions, we can conclude from this that:

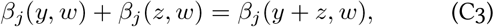

with *w* = *ψ_j_* (**x**, **a**, x_*j*_ = *y* + *z*, x_*k*_ = 0). Therefore *β* must be proportional in its first argument, i.e. the right-hand side has the shape:

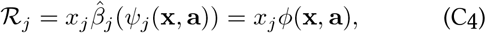

with some impact function 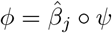.

## Appendix D: Inconsistencies in the Model from Ref. 16

We here provide explicit examples for an inconsistent behavior of the model described by Eq. 20.

First, in Fig. 5 we show a simulation similar to Fig. 1 exhibiting different results when simulating the same scenario in two different ways. This simulation also exposes another problem of the model, namely that it contradicts the underlying assumption that a population also represent its footprint. Without dilution, this footprint cannot decrease and thus populations cannot decrease, which however happens in the simulation in Fig. 5.

Second, in the following we provide an arithmetic example that features no growth interaction and thus does not rely on how we chose identical populations to affect each other’s growth (see Appendix F) – a choice that is not clear without experiment. We consider the case of three populations {1, 2, 3} =: *J* with the first two populations having identical properties. We choose *g_j_* = 1 ∀*j* ∈ *J*, *a_j,k_* = 0 ∀*j,k* ∈ *J* × *J*, i.e., the growth term is not affected by interaction and is always 1. Finally, we let the coefficients of the capacity term be:

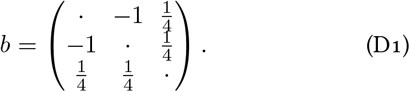

*b*_12_ = –1 and *b*_21_ = –1 reflects that two populations with identical properties deplete each other’s niches. Now, consider two states of the community 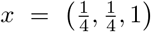 and 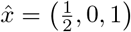. As the first two populations are indistinguishable, these states describe an equivalent situation. Thus, they should also evolve equivalently, i.e., the temporal derivative of the summed populations 1 and 2 should be the same in both cases: 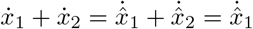. However,

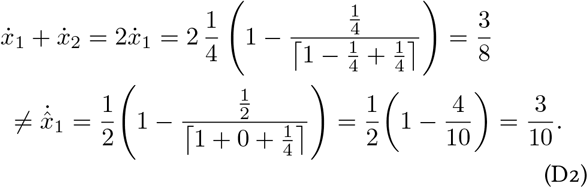

## Appendix E: Deriving a New Model for UTI strains – the Legwork

Inserting our ansatz (Eq. 23) into our first requirement (Eq. 16), we obtain:

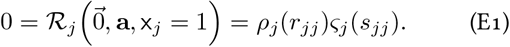

Assuming that the two factors do not “take turns” in being zero for different *j*, this means that either *ς_j_* (*j*) = 0 or *ς_j_*(*s_jj_*) = 0. Without loss of generality, we assume that the latter applies, thus assigning *ς_j_* the role of quantifying the carrying capacity. Furthermore, we choose *s_jj_* = 1 and *ς_j_* (0) = 1. These are normalization choices, as they can be compensated by including a respective factor in *ς_j_* or *ρ_j_* respectively. Using this and expanding Eq. 18, we obtain:

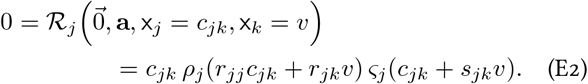

Assuming that *ς_j_* is again responsible for the product being zero and it has only one root, namely 1, we arrive at: *c_jk_* + *s_jk_ υ* = 1, and thus: 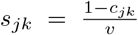. Note that since *s_jj_* = 1, this is consistent with our choice of *c_jj_* = 1 – *υ* (see Appendix F).

Using the above, we can expand Eqs. 15 and 17:

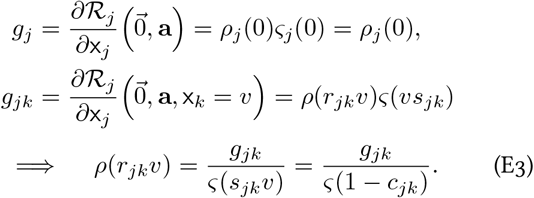

We choose the arguably simplest function to fulfill the criteria for *ρ*, namely *ρ_j_*(*z*)≔ *g_j_* + *z*. This has the consequence:

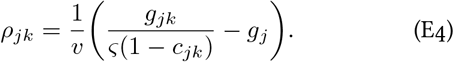

A group of functions fulfilling the criteria for *ς* is: *ς_j_* (*z*) ≔ 1 – ⌈*z*⌉^*q*^ with *q* > 0 and ⌈*z*⌉ ≔ max(0, *z*). Here, the free parameter *q* controls how early and smoothly the saturation effect of a occupied niche kicks in. Note that this choice results in terms similar to what Ref. 69 named *hyperlogistic*.

Finally, like Ref. 16, we constrain the growth and capacity term to be non-negative to avoid the occasional implausible result. For example, we do not allow negative growth because we equate the abundance of a population with its footprint, which cannot be undone, and we lack the data to capture cell death. Putting everything together, we arrive at the model:

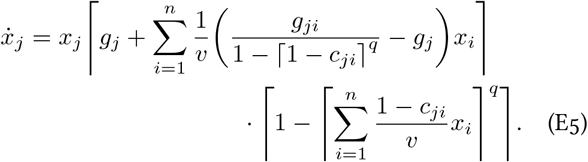

A problem with this model is that for 0 < *x_k_* < 1, we have: 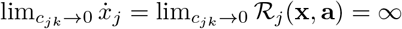. Now, *c_jk_* = 0 means that there is no growth of strain *j* in the medium conditioned by strain *k* and thus we already have a problem with experimentally determining *g_jk_*. Thus, one might argue that the actual point of the singularity requires a dedicated case distinction anyway. However, 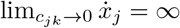. also means that 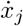 becomes arbitrarily large for small *c_jk_*. A way to address this problem is to consider the case *q* → ∞, or more specifically:

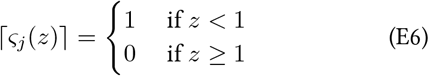

In this case, the term *ς*(1 – *c_jk_*) in Eq. E3 can be assumed to be 1 (otherwise, we would have the aforementioned problem of not being able to experimentally determine *g_jk_*). This eliminates the singularity, but also renders the model not continuously differentiable.

In our simulations, we therefore make a trade-off between complying with Eq. 17 and the numerical benefits of a continuously differentiable model by setting *q* =10 and approximating *ς*(1 – *c_jk_*) ≈ lim_*p*→∞_ *ς*(1 – *c_jk_*) = 1 in Eq. E3, thus arriving at:

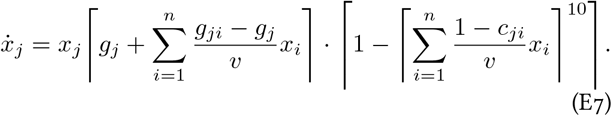

## Appendix F: Comparing the Old and New UTI Model with Experimental Data

To test the applicability of our UTI model (Eq: 24) in practice, we compared its predictions of in-vitro experiments with known outcome to those of the previous model (Eq. 20).

In the in-vitro experiments from Ref. 16 (cf. Fig. S7 B therein), each strain was inoculated at a fixed optical density (OD_600_ = 0.001) grown for 4 × 30 h and diluted by a factor of 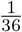 in between. Each experiment was performed in triplicate. Final abundances were determined from colony-forming units on Chromagar.

We mimicked this experimental procedure in simulations using both models. For these, we use the data from Ref. 16 as is, with the exception of data describing interactions between two identical strains: First, in the experiment, a strain cannot grow on the portion of the medium that is its own supernatant, but only on the portion that is fresh medium, which makes up 1 – *υ* = 0.6 of the medium. We set *c_ii_* = 1 – *υ* to adhere to this ideal. Second, in the medium conditioned by itself, a strain’s growth rate should at best slightly lower than in an unconditioned medium and at worst be proportional to the concentration of nutrients, and thus to 1 – *υ*. We therefore restrict *g_ii_* to the interval [(1 – *υ*)*g_i_*, (1 – *ϵ*)*g_i_*] with *ε* = 0.01. Without these adjustments, we would obtain implausible results, e.g., in case of *c_ii_* > 1 – *υ*, the respective strain could never stop growing since it effectively increases the size of its own niche.

We performed all simulations with JiTCODE [70] using the DoPri5 method. To obtain continuity as required by the integration method, we approximate 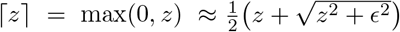 with *ϵ* = 0.001.

We converted abundances in the simulation results to optical densities by undoing the respective normalization of abundances (fixing *c_j_* = 1, see Fig. 4). We then converted the optical densities to displayed abundances (in Figs. 5 and 6) by approximating that optical density is proportional to biovolume.

**FIG. 6.**
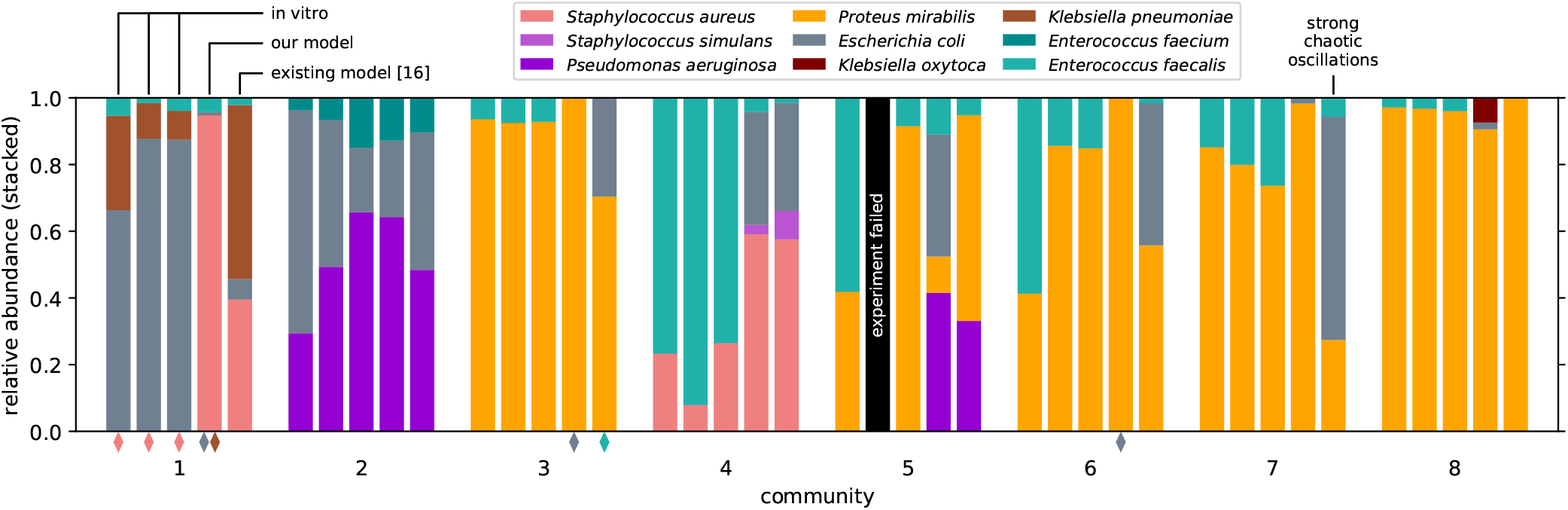
Comparison of the final relative abundances of in-vitro experiments from Ref. 16 (Fig. S7 B) and simulations with our model (Eq. 24) and the model from Ref. 16 (Eq. 20). Communities are numbered as in Ref. 16. Diamonds at the bottom indicate an abundance of the respective population between 10^−4^ and 10^−2^. Community 2 featured two strains of *Enterococcus faecium*, for which we only report the summed populations since they could not be distinguished in experiment. All simulations converged, except for the model from Ref. 16 and Community 7, which exhibited strong chaotic fluctuations.

The results of the simulation are shown in Fig. 6. Note that quantitatively predicting the population dynamics in experimental scenarios like this without in-depth knowledge about the involved microbes is a highly difficult challenge. Moreover, the high-throughput interaction data used to build the models is restricted; for example, it does not feature higher-order interactions, and the supernatant used to determine interactions will not contain toxins whose production is triggered by products of their target. Therefore, neither model can be expected to make perfect predictions.

We find that both models are in equally good or bad agreement with the experiment for six communities (1, 2, 4, 5, 6, and 8), while the predictions of our model (Eq. 24) are better for two communities (3 and 7). Given the low number of samples, we refrain from further quantifying the agreements. These results indicate that models satisfying our consistency criteria are at least equally suitable for describing community dynamics. Our results do not challenge the conclusions of Ref. 16, as both models yield similar results with respect to the key findings, in particular the stability of steady states. This is plausible since both models have the same fixed points if the growth term is ignored and *c_jk_* < 1 ∀*j, k* and can thus be expected to yield similar final states. Thus differences between the models primarily arise when the system is diluted before it has equilibrated, which explains why the modelled communities tend to differ little (2, 4, and 8) or strongly (1, 5, 6, and 7).

