## Supplementary material for "Building clone-consistent ecosystem models": Code for Simulations: colours.py

```
species_colours = {
		'Staphylococcus aureus' : "lightcoral",
		'Staphylococcus haemolyticus' : "mediumorchid",
		'Staphylococcus simulans' : "mediumorchid",
		'Proteus mirabilis' : "orange",
		'Enterobacter cloacae' : "black",
		'Enterococcus faecium' : "darkcyan",
		'Enterococcus faecalis' : "lightseagreen",
		'Escherichia coli' : "slategrey",
		'Pseudomonas aeruginosa' : "darkviolet",
		'Klebsiella pneumoniae' : "sienna",
		'Klebsiella oxytoca' : "maroon",
}
```
