## Supplementary material for "Building clone-consistent ecosystem models": Code for Simulations: Coutlee_Inconsistency.ipynb

```
# %% [code]
from matplotlib.pyplot import subplots
import numpy as np

# %% [markdown]
The following class implements the Coutlee model with the possibility to clone one population (and halve the initial abundance).

# %% [code]
class CoutleeModel(object):
    def __init__(self,α,β,initial,times,populations,colors):
        assert len(α) == len(β) == len(initial) == len(populations) == len(colors)
        self.α = α
        self.β = β
        self.initial = initial
        self.times = times
        self.populations = populations
        self.colors = colors
        self.clones = []
    
    def make_clone(self,target):
        assert not self.clones
        N = len(self.α)
        self.clones = (target,N)

        self.α = np.array([*self.α,self.α[target]])

        new_β = np.empty((N+1,N+1))
        new_β[:N,:N] = self.β
        for i in range(N):
            new_β[i,N] = self.β[i,target]
            new_β[N,i] = self.β[target,i]
        new_β[N,N] = self.β[target,target]
        self.β = new_β

        self.initial = np.array([*self.initial,self.initial[target]])
        self.initial[[target,N]] /= 2
    
    def simulate(self):
        from jitcode import jitcode, y as n
        from symengine import log
        N = len(self.α)
        f = [
            n(i)*( self.α[i] + sum( self.β[i,j]*log(n(j)) for j in range(N) ) )
            for i in range(N)
        ]
        ODE = jitcode( f, verbose=False )
        ODE.set_integrator("RK45")
        ODE.set_initial_value(self.initial)

        return np.vstack([ ODE.integrate(time) for time in times ]).T
    
    def plot(self,axes,**kwargs):
        """
        Plot a timeseries for the given scenario on the given axes, summing up the dynamical variables in `clones`.
        """
        results = self.simulate()
        
        lines = []
        for (i,result) in enumerate(results):
            if self.clones:
                # Sum up the result for the cloned populations in one of them and ignore the other one.
                if i==self.clones[0]:
                    result += results[self.clones[1]]
                elif i==self.clones[1]:
                    continue
            lines.extend(axes.plot(
                self.times, result,
                color=colors[i], label=populations[i],
                **kwargs
            ))
        return lines

# %% [markdown]
Here we define the specific parameters, initial conditions, and time span we want to simulate:

# %% [code]
α = np.array([0.3,-0.3])
β = np.array([
    [0.5,-1  ],
    [0.8,-0.5],
])
N = len(α)
initial = np.array([2.0,6.0])
times = np.linspace(0,22,1000)

populations = ["prey","predator"]
colors = ["blue","orange"]

# %% [markdown]
Finally, we perform the plot:

# %% [code]
model = CoutleeModel(α,β,initial,times,populations,colors)

fig,axes = subplots(figsize=(6,2))
fig.patch.set_alpha(0)

styles = { "plain":"-", "with clone":"--" }
from matplotlib.lines import Line2D
lines = model.plot(axes,linestyle=styles["plain"])
model.make_clone(target=1)
model.plot(axes,linestyle=styles["with clone"])

dummy_lines = [
    axes.plot( [],[], color="black", linestyle=style, label=name )[0]
    for name,style in styles.items()
]

for lines,loc in [ (dummy_lines,"upper left"), (lines,"upper right") ]:
    legend = axes.legend( handles=lines, loc=loc, prop={"size":8} )
    axes.add_artist(legend)
    
import matplotlib.ticker as ticker
axes.set_xlim(times[0],times[-1])
axes.set_xlabel("time")
axes.xaxis.set_major_locator(ticker.MultipleLocator(7))
    
axes.set_ylabel("abundance $x_i$")
axes.set_ylim(ymin=0,ymax=11)
axes.yaxis.set_tick_params(right=True)
axes.set_yticks([0,2,6,10]);
```
