## Supplementary material for "Building clone-consistent ecosystem models": Code for Simulations: in_vitro.py

```
experimental_data = {
		('in vitro, replicate 1', 0): {
				'Klebsiella pneumoniae': 195,
				'Enterococcus faecalis': 37,
				'Escherichia coli': 456,
				'Staphylococcus aureus': 1,
			},
		('in vitro, replicate 2', 0): {
				'Klebsiella pneumoniae': 76,
				'Enterococcus faecalis': 11,
				'Escherichia coli': 622,
				'Staphylococcus aureus': 1,
			},
		('in vitro, replicate 3', 0): {
				'Klebsiella pneumoniae': 49,
				'Enterococcus faecalis': 22,
				'Escherichia coli': 494,
				'Staphylococcus aureus': 1,
			},
		('in vitro, replicate 1', 1): {
				'Enterococcus faecium': 28,
				'Escherichia coli': 500,
				'Pseudomonas aeruginosa': 220,
			},
		('in vitro, replicate 2', 1): {
				'Enterococcus faecium': 43,
				'Escherichia coli': 286,
				'Pseudomonas aeruginosa': 320,
			},
		('in vitro, replicate 3', 1): {
				'Enterococcus faecium': 115,
				'Escherichia coli': 148,
				'Pseudomonas aeruginosa': 504,
			},
		('in vitro, replicate 1', 2): {
				'Enterococcus faecalis': 71,
				'Proteus mirabilis': 1040,
			},
		('in vitro, replicate 2', 2): {
				'Enterococcus faecalis': 90,
				'Proteus mirabilis': 1112,
			},
		('in vitro, replicate 3', 2): {
				'Enterococcus faecalis': 80,
				'Proteus mirabilis': 1032,
			},
		('in vitro, replicate 1', 3): {
				'Enterococcus faecalis': 23,
				'Staphylococcus aureus': 7,
			},
		('in vitro, replicate 2', 3): {
				'Enterococcus faecalis': 35,
				'Staphylococcus aureus': 3,
			},
		('in vitro, replicate 3', 3): {
				'Enterococcus faecalis': 25,
				'Staphylococcus aureus': 9,
			},
		('in vitro, replicate 1', 4): {
				'Enterococcus faecalis': 118,
				'Proteus mirabilis': 85,
			},
		('in vitro, replicate 2', 4): {
				#'Enterococcus faecalis': 262,
			},
		('in vitro, replicate 3', 4): {
				'Enterococcus faecalis': 148,
				'Proteus mirabilis': 1600,
			},
		('in vitro, replicate 1', 5): {
				'Enterococcus faecalis': 128,
				'Proteus mirabilis': 90,
			},
		('in vitro, replicate 2', 5): {
				'Enterococcus faecalis': 127,
				'Proteus mirabilis': 756,
			},
		('in vitro, replicate 3', 5): {
				'Enterococcus faecalis': 136,
				'Proteus mirabilis': 764,
			},
		('in vitro, replicate 1', 6): {
				'Enterococcus faecalis': 232,
				'Proteus mirabilis': 1336,
			},
		('in vitro, replicate 2', 6): {
				'Enterococcus faecalis': 200,
				'Proteus mirabilis': 796,
			},
		('in vitro, replicate 3', 6): {
				'Enterococcus faecalis': 180,
				'Proteus mirabilis': 504,
			},
		('in vitro, replicate 1', 7): {
				'Enterococcus faecalis': 40,
				'Proteus mirabilis': 1352,
			},
		('in vitro, replicate 2', 7): {
				'Enterococcus faecalis': 47,
				'Proteus mirabilis': 1408,
			},
		('in vitro, replicate 3', 7): {
				'Enterococcus faecalis': 53,
				'Proteus mirabilis': 1280,
			},
	}
```
