## Supplementary material for "Building clone-consistent ecosystem models": Code for Simulations: loaddata.py

```
from scipy.io import loadmat
import numpy as np

with open("communities.csv","r") as datei:
	lines = [ line.strip().split("\t") for line in datei]

communities = []
names = {}

while lines:
	assert len(lines[0]) == 3
	comm_id = lines[0].pop(2)
	assert comm_id.startswith("Comm")
	assert int(comm_id.partition("Comm")[2]) == len(communities)+1
	
	community = []
	while lines and len(lines[0])==2:
		strain = lines.pop(0)
		name = strain[0]
		try:
			number = int(strain[1])
		except ValueError:
			if __name__ == "__main__":
				print(f"Could not parse {strain}")
			continue
		names[number] = name
		community.append(number)
	
	communities.append(community)

order = [56, 2, 91, 42, 59, 53, 10, 62, 80, 34, 76, 22, 18, 19, 16, 45, 68, 36, 74, 79, 4, 44, 23, 9, 5, 21, 30, 51, 87, 41, 58, 92, 54, 72, 83, 37, 25, 69, 60, 84, 78, 63, 13, 82, 90, 1, 7, 33, 64, 14, 47, 11, 66, 6, 31, 71, 35, 20, 3, 61, 28, 81, 88, 38, 46, 43, 65, 55, 15, 50, 75, 77]

growths = {
		i+1: growth
		for i,growth in enumerate(loadmat("mean_GR_spentHRs.mat")["mean_GR_spentHRs"][0])
	}

ODs = {
		i+1: OD
		for i, OD in enumerate(loadmat("mean_maxOD_spentHRs.mat")["mean_maxOD_spentHRs"][0])
	}

# second index is the conditioning species
conditioned_growths = {
		(order[j],order[i]) : conditioned_growth
		for i,line in enumerate(loadmat("GR_3.mat")["GR_Int_spentS_reorder_HR_cleaned_20160514_3"])
		for j,conditioned_growth in enumerate(line)
	}

# second index is the conditioning species
conditioned_yields = {
		(order[j],order[i]) : conditioned_yield
		for i,line in enumerate(loadmat("MaxOD_3.mat")["MaxOD_Int_spentS_reorder_20160514_3"])
		for j,conditioned_yield in enumerate(line)
	}

# second index is the conditioning species
denormalised_growths = {
		(i,j): conditioned_growths[i,j]*growths[i]
		for i,j in conditioned_growths.keys()
	}

volumes = {}
with open("species_sizes.csv","r") as datei:
	for line in datei:
		name,volume,source = line.strip().split("\t")
		volumes[name] = float(volume)

def get_community(strain):
	for i,community in enumerate(communities):
		if strain in community:
			return i+1
	else:
		raise AssertionError

def get_genus(strain):
	return strain.split()[0]
```
