## Supplementary material for "Building clone-consistent ecosystem models": Code for Simulations: Model_Comparison.ipynb

```
# %% [code]
from matplotlib.pyplot import subplots
import numpy as np
from loaddata import communities, names
from UTI_simulator import NewModel,OldModel
from colours import species_colours

# %% [markdown]
The parameters of the experimental scenario:

# %% [code]
transfer_number = 4
transfer_distance = 30*60
transfer_dilution = 1/36

# Only use the first eight communities (because only those were measured experimentally):
communities = communities[:8]

# %% [markdown]
Load the experimental data and convert it to ratios:

# %% [code]
from in_vitro import experimental_data

def convert_to_ratios(experiment):
    total = sum(experiment.values())
    if total:
        for key in experiment:
            experiment[key] /= total
    else:
        experiment["missing"] = 1

def convert_all_to_ratios(dataset):
    for experiment in dataset.values():
        convert_to_ratios(experiment)
    return dataset

ratios = convert_all_to_ratios(experimental_data)

# %% [markdown]
Perform the simulations and join them with the experimental results in `ratios`:

# %% [code]
def get_ratios(community, Model):
    """
    Returns the final ratios of abundances (in CFUs) in form of a dictionary.
    """
    ODE = Model(community)
    ODE.set_initial_value_by_OD( np.full(len(community),0.001) )
    
    for _ in range(transfer_number):
        state = ODE.integrate(ODE.t+transfer_distance)
        ODE.dilute(transfer_dilution)
    
    normalised_state = state/sum(state)
    
    result = {}
    for i,strain in enumerate(community):
        species = names[strain]
        if normalised_state[i]:
            try:
                result[species] += normalised_state[i]
            except KeyError:
                result[species] = normalised_state[i]
    return result

for name,Model in [ ("our model",NewModel), ("existing model [16]",OldModel) ]:
    for number,community in enumerate(communities):
        ratios[name,number] = get_ratios(community,Model)

# %% [markdown]
Organize our species and experiments:

# %% [code]
# Using a dictionary (instead of a set) to preserve insertion order
all_experiments = {key[0]:None for key in ratios.keys()}

all_species = {species for experiment in ratios.values() for species in experiment}
# because they doen’t survive anyway (otherwise you would see gaps in the bar plot):
all_species.remove("Enterobacter cloacae")
all_species.remove("Staphylococcus haemolyticus")

# Selected order for better readability of the plot:
plotted_species = [
    'Staphylococcus aureus',
    'Staphylococcus simulans',
    'Pseudomonas aeruginosa',
    'Proteus mirabilis',
    'Escherichia coli',
    'Klebsiella oxytoca',
    'Klebsiella pneumoniae',
    'Enterococcus faecium',
    'Enterococcus faecalis',
]

assert set(plotted_species)|{"missing"} == all_species

# %% [markdown]
Plot everything:

# %% [code]
n = len(communities) 
# number of bar spaces per community (number of experiments + gap):
m = len(all_experiments)+1 

species_colours["missing"] = "black"

fig,axes = subplots(figsize=(13,3))
fig.patch.set_alpha(0)

# The main bar plots:
current_bar_heights = np.zeros(m*n)
bar_positions = np.arange(1,n+1,1/m)-0.5+1/m

for species in plotted_species+["missing"]:
    added_bars = np.zeros_like(current_bar_heights)
    for i,experiment in enumerate(all_experiments):
        for j in range(n):
            try:
                added_bars[j*m+i] = ratios[experiment,j][species]
            except KeyError:
                pass

    axes.bar(
            bar_positions, added_bars, bottom=current_bar_heights,
            label = species if species!="missing" else "__nolegend__",
            width = 0.8/m,
            color = species_colours[species],
        )
    current_bar_heights += added_bars

# Diamonds under plot for abundances below 1e−2 and above 1e−4:
for i,experiment in enumerate(all_experiments):
    for j in range(n):
        dot_colours = []
        for species in plotted_species:
            try:
                result = ratios[experiment,j][species]
            except KeyError:
                pass
            else:
                if 1e-4<result<1e-2:
                    dot_colours.append(species_colours[species])

        k = len(dot_colours)
        Δ = 0.03
        offsets = np.linspace(-(k-1)*Δ,(k-1)*Δ,k)
        for dot_colour,offset in zip(dot_colours,offsets):
            axes.text(
                    bar_positions[j*m+i]+offset, -0.05, "⧫",
                    horizontalalignment = 'center',
                    color = dot_colour,
                )


# legend for the different experiments/simulations
annotation_style = {
    "arrowprops": { "arrowstyle":"-", "connectionstyle":"angle, angleA=0, angleB=90" },
    "horizontalalignment": "left",
    "fontsize": 8,
}

for i,(bar_position,experiment) in enumerate(zip(bar_positions,all_experiments)):
    vskip = 0.09
    if i<2:
        text = " "
        i = 2
    else:
        text = experiment.split(", replicate")[0]
    axes.annotate( text, (bar_position,1), (1.5,0.88+vskip*m-vskip*i), **annotation_style )

# Marking the incident of chaos:
index = next(
    i for i,name in enumerate(all_experiments)
    if name.startswith("model from de Vos")
)
hpos = 6.5+(index+1)/m
axes.annotate(
        "strong\nchaotic\noscillations",
        (hpos,1), (hpos,1.1),
        **annotation_style,
        ha = "center"
    )

# Marking the failed experiment:
for i,experiment in enumerate(all_experiments):
    for j in range(n):
        if "missing" in ratios[experiment,j]:
            axes.text(
                bar_positions[j*m+i], 0.5,
                "experiment failed",
                rotation=90, rotation_mode='anchor',
                ha="center", va="center",
                fontsize=8, color="white",
            )

# Legends, etc.:
axes.legend(
    bbox_to_anchor = (0.29,1.03),
    ncol = 3, columnspacing = 1,
    prop = {'style':'italic','size':8}
)

axes.set_xlabel("community")
axes.set_ylabel("relative abundance (stacked)")
axes.set_xlim( bar_positions[0]-1.4/m, bar_positions[-1]+0.4/m )
axes.yaxis.set_tick_params(right=True)
axes.xaxis.set_tick_params(pad=10, bottom=False)

# %% [code]
fig.savefig("simulations.pdf",bbox_inches="tight")

# %% [code]
```
