## Supplementary material for "Building clone-consistent ecosystem models": Code for Simulations: UTI_Inconsistency.ipynb

```
# %% [code]
from matplotlib.pyplot import subplots
import numpy as np
from loaddata import communities, names, get_genus
from UTI_simulator import OldModel
from colours import species_colours

# %% [markdown]
First we select a community and clone a selected member (here the zeroth one).
It’s easier to distinguish scenarios by initial conditions only, since a population starting at zero has no impact anyway.
We also choose initial conditions and the total simulation time.

# %% [code]
community = [34,36,35,37] # reordering to make the legend smaller
assert sorted(community)==communities[2]

clone_target = 0
clones = [clone_target,len(community)]
community.append(community[clone_target])

max_t = 1200

base_initial = [ 0.029, 0.03, 0.7, 0.1, 0 ]

# %% [markdown]
Here we set up the case with clones as well as with perturbed initial conditions:

# %% [code]
cloned_initial = np.copy(base_initial)
cloned_initial[clones[1]] = cloned_initial[clones[0]]
cloned_initial[clones] /= 2

amp = 0.01
perturbation = np.random.choice([1-amp,1+amp],size=5)
perturbed_initial = base_initial*perturbation

scenarios = [
    { "name": "plain"     , "initial":      base_initial, "style": "-"  },
    { "name": "with clone", "initial":    cloned_initial, "style": "--" },
    { "name": "perturbed" , "initial": perturbed_initial, "style": ":"  },
]

# %% [code]
ODE = OldModel(community)

# %% [markdown]
Finally, we run and plot everything:

# %% [code]
fig,axes = subplots(figsize=(6,3))
fig.patch.set_alpha(0)

for scenario in scenarios:
    ODE.set_initial_value(scenario["initial"],0)
    times = np.linspace(0,max_t,1000)
    results = np.vstack([ODE.integrate(time) for time in times]).T
    
    # Summing the abundances of the cloned populations into one:
    results[clones[0]] += results[clones[1]]
    
    for strain,result in zip(community[:-1],results[:-1]):
        axes.plot(
            times/60, result,
            color = species_colours[names[strain]],
            linestyle = scenario["style"],
        )

import matplotlib.ticker as ticker
axes.set_xlim(0,max_t/60)
axes.set_xlabel("time in hours")
axes.xaxis.set_major_locator(ticker.MultipleLocator(5))

axes.set_ylabel("abundance (a.u.)")
axes.yaxis.set_tick_params(right=True)
axes.set_yscale("log")

lines = [
    axes.plot([],[],color="black",linestyle=scenario["style"],label=scenario["name"])[0]
    for scenario in scenarios
]
scenario_legend = axes.legend(handles=lines,loc="lower left",bbox_to_anchor=(0,1.02))
axes.add_artist(scenario_legend)

lines = [
    axes.plot([],[],color=species_colours[names[strain]],label=get_genus(names[strain]))[0]
    for strain in community[:-1]
]
axes.legend(
        handles = lines,
        prop = {'style':'italic'},
        loc = "lower right",
        bbox_to_anchor = (1,1.02),
        ncol = 2,
    )
```
