## Supplementary material for "Building clone-consistent ecosystem models": Code for Simulations: UTI_simulator.py

```
import numpy as np
from sympy import IndexedBase, Piecewise, Rational, Symbol, sqrt
from jitcode import jitcode, y
from loaddata import growths, denormalised_growths, conditioned_yields, names, ODs, volumes, get_genus

n = Symbol("n",integer=True)

x = IndexedBase("x")
Y = IndexedBase("Y")
g = IndexedBase("g")

v = Rational(4,10)
ε = 0.001

# The function z ↦ max(0,z) = ⌊z⌋ ≈ ½(z + √(z²+ε²))
minzero = lambda z: (z + sqrt(z**2+ε**2))/2

class UTI_simulator(jitcode):
	"""
	The base class for UTI simulations.
	This needs to be inherited from and equipped with a method `model`.
	"""
	def __init__(self,community):
		self.community = community
		self.helper_index = 0
		
		# Conversion from relative ODs to CFUs:
		self.correction = np.array([ODs[i]/volumes[get_genus(names[i])] for i in community])
		
		substitutions = self.substitutions()
		
		# Convention:
		# uppercase: strain IDs according to database
		# lowercase: dynamical variable IDs
		
		f = []
		helpers = []
		for i,I in enumerate(self.community):
			model_i,helpers_i = self.model(i,I)
			f.append(model_i.subs(substitutions))
			for helper in helpers_i:
				helpers.append((helper[0],helper[1].subs(substitutions)))
		
		super().__init__(f,helpers=helpers,verbose=False)
		self.generate_f_C(simplify=False)
		self.set_integrator("RK45")
	
	def get_helper(self):
		"""
		This method exists to avoid redundant helpers.
		"""
		self.helper_index += 1
		return Symbol(f"helper{self.helper_index}")
	
	def substitutions(self):
		"""
		Returns a dictionary that translates the abstract model formulation to numbers or symbols used by the integrator.
		"""
		subs = { x[j]:y(j) for j,J in enumerate(self.community) }
		for J in self.community:
			subs[g[J]] = growths[J]
			for K in self.community:
				subs[Y[J,K]] = conditioned_yields[J,K]
				subs[g[J,K]] = denormalised_growths[J,K]
			subs[Y[J,J]] = 1-v
			subs[g[J,J]] = min(growths[J]*0.99,subs[g[J,J]])
			subs[g[J,J]] = max(growths[J]*(1-v),subs[g[J,J]])
		return subs
	
	def integrate(self,t):
		return super().integrate(t)*self.correction
	
	def dilute(self,dilution_factor):
		super().set_initial_value(self.y*dilution_factor,self.t)
	
	def set_initial_value(self,initial,*args):
		super().set_initial_value(initial/self.correction,*args)
	
	def set_initial_value_by_OD(self,initial_ODs,*args):
		initial_values = [
				initial_OD/ODs[strain]
				for strain,initial_OD in zip(self.community,initial_ODs)
			]
		super().set_initial_value(initial_values,*args)

a = lambda J,K: g[J,K]/g[J]-1
b = lambda J,K: Piecewise( (Y[J,K]-1,Y[J,K]>1), ((Y[J,K]-1)/v,True) )
class OldModel(UTI_simulator):
	def model(self,j,J):
		growth_sum = self.get_helper()
		yield_sum = self.get_helper()
		helpers = [
				(growth_sum, 1 + sum( a(J,K)*x[k] for k,K in enumerate(self.community) if k!=j ) ),
				(yield_sum , 1 + sum( b(J,K)*x[k] for k,K in enumerate(self.community) if k!=j ) ),
			]
		model = x[j] * g[J] * minzero(growth_sum) * (1-x[j]/minzero(yield_sum))
		return model, helpers

q = 10
class NewModel(UTI_simulator):
	def model(self,j,J):
		growth_term = self.get_helper()
		yield_sum = self.get_helper()
		helpers = [
				( growth_term, g[J] + sum( (g[J,K]-g[J])/v*x[k] for k,K in enumerate(self.community) ) ),
				( yield_sum, sum( (1-Y[J,K])/v*x[k] for k,K in enumerate(self.community) ) ),
			]
		model = x[j] * minzero(growth_term) * minzero(1-minzero(yield_sum)**q)
		return model, helpers
```
